# AAVs Targeting Human Carbonic Anhydrase IV Enhance Gene Delivery to the Brain

**DOI:** 10.1101/2025.04.21.649868

**Authors:** Changfan Lin, Xinhong Chen, Jonathan D. Hoang, Fiona Ristic, Yujie Fan, Seongmin Jang, Jin Hyung Alex Chung, Erin E. Sullivan, Tomasz Gawda, Bill Kavvathas, Irene Tran, Yitong Li, Andrew D. Steele, Timothy F. Shay, Viviana Gradinaru

## Abstract

Clinically approved gene therapies based on natural adeno-associated virus (AAV) serotypes have restricted applications, particularly in the brain, due to their poor targeting, high dose requirements, and resulting safety concerns. Directed evolution of enhanced AAV capsids in mice or non-human primates (NHPs) has resulted in markedly improved performance in those species, but inter-species differences present a serious challenge for translating these vectors into human therapies. Here, we engineer AAVs to target human carbonic anhydrase IV (CA-IV), a recently identified blood-brain barrier (BBB) transcytosis receptor. Among known transcytosis receptors, CA-IV is notable for its relatively specific expression in brain endothelial cells and the potency of AAVs that target it in mice. CA-IV’s AAV binding site, and thus the mouse vectors’ enhanced brain potency, is not conserved across species, so we employed a two-phase engineering strategy to identify AAVs optimized for human CA-IV-dependent gene delivery to the brain. We first used in vitro receptor-based selection of a vast AAV library to exclude capsids that do not bind human CA-IV, followed by in vivo selection in “humanized” mice expressing human CA-IV in brain endothelial cells. Notably, we find that human CA-IV binding capsid variants that were poorly enriched in the pull-down selection outperform strong binders in vivo. The most promising vector, AAV-hCA4-IV77, engages human CA-IV to achieve 100-fold greater brain transduction than AAV9, with robust neuronal and astrocytic coverage throughout multiple brain regions. These results advance our understanding of receptor-targeted capsid design and support the therapeutic potential of human CA-IV-engaging AAVs.

## Introduction

Gene therapy is a rapidly advancing field that treats genetic disorders by addressing their molecular origins. Adeno-associated viral vectors (AAV) are preferred gene delivery vehicles due to their favorable safety profile compared to other viral vectors and their lack of known pathogenicity in humans^1–3^. Currently, there are over 300 clinical trials exploring AAV-based therapies, and five treatments based on naturally evolved AAV serotypes have already received approval from the US Food and Drug Administration (FDA)^4–9^. These include Zolgensma, which utilizes the AAV9 capsid for intravenous targeting of the nervous system to treat spinal muscular atrophy in children up to two years of age^10,11^. While gene therapies utilizing natural AAV serotypes are dramatically changing the trajectory of select genetic disorders, many cell types and tissues are transduced only at high systemic doses, negatively impacting the safety and efficacy of existing treatments, and precluding application to some indications^12–15^. Thus, for AAV-based gene therapy to reach its full potential, further optimized AAV vectors are required.

Recent advances in engineering AAV capsids for systemic delivery through *in vivo* directed evolution have markedly enhanced specificity and potency in mice and non-human primates (NHPs)^16–23^. This engineering approach involves introducing a diversified library of AAV capsid variants into an animal and then recovering those that effectively reached the target tissues or cell types. The resulting engineered systemic AAVs have been broadly applied as research tools, which has revealed that the capsids’ properties are often specific to the model organism in which they were developed, with tropism changing or disappearing in different host genetic contexts^24–26^. This limitation is primarily due to receptor differences between species. Performing directed evolution in NHPs increases the translational potential of engineered capsids, but is resource-intensive and does not fully eliminate the risk of altered behavior when applied to humans. Thus, a different approach for engineering human-optimized systemic AAVs is needed.

The central nervous system is an especially challenging target for gene therapy due to the blood-brain barrier (BBB). This protective structure, composed primarily of tightly joined endothelial cells, restricts the entry of most therapeutic agents into the brain^27–31^. Recent efforts by our lab and others have identified the BBB receptors utilized by certain engineered systemic capsids to cross into the brain through receptor-mediated transcytosis^32–36^. One such receptor is carbonic anhydrase IV (CA-IV), which facilitates the efficient BBB crossing of AAV variants 9P31 and 9P36 in mice^18,32^. Importantly, CA-IV is also present in NHP and human BBB endothelial cells (Figure S1A-B)^37,38^ and exhibits a more CNS-focused expression pattern than other established BBB receptors, such as low-density lipoprotein receptor-related protein 6 (LRP6), transferrin receptor (TfR1) and insulin receptor (INSR) (Figure S1C). This restricted distribution may offer advantages for targeted drug delivery to the brain, potentially reducing off-target effects in other tissues. While CA-IV homologs exhibit significant sequence conservation across species, they are not identical. For instance, rhesus macaque and marmoset CA-IV share 85% amino acid identity with human CA-IV, whereas mouse CA-IV shares only 55% identity (Figure 1A). As a result, AAVs 9P31 and 9P36 bind exclusively to mouse, but not primate, CA-IV, with 9P31 crossing the BBB similarly to AAV9 in macaque^39^.

**Figure 1:**
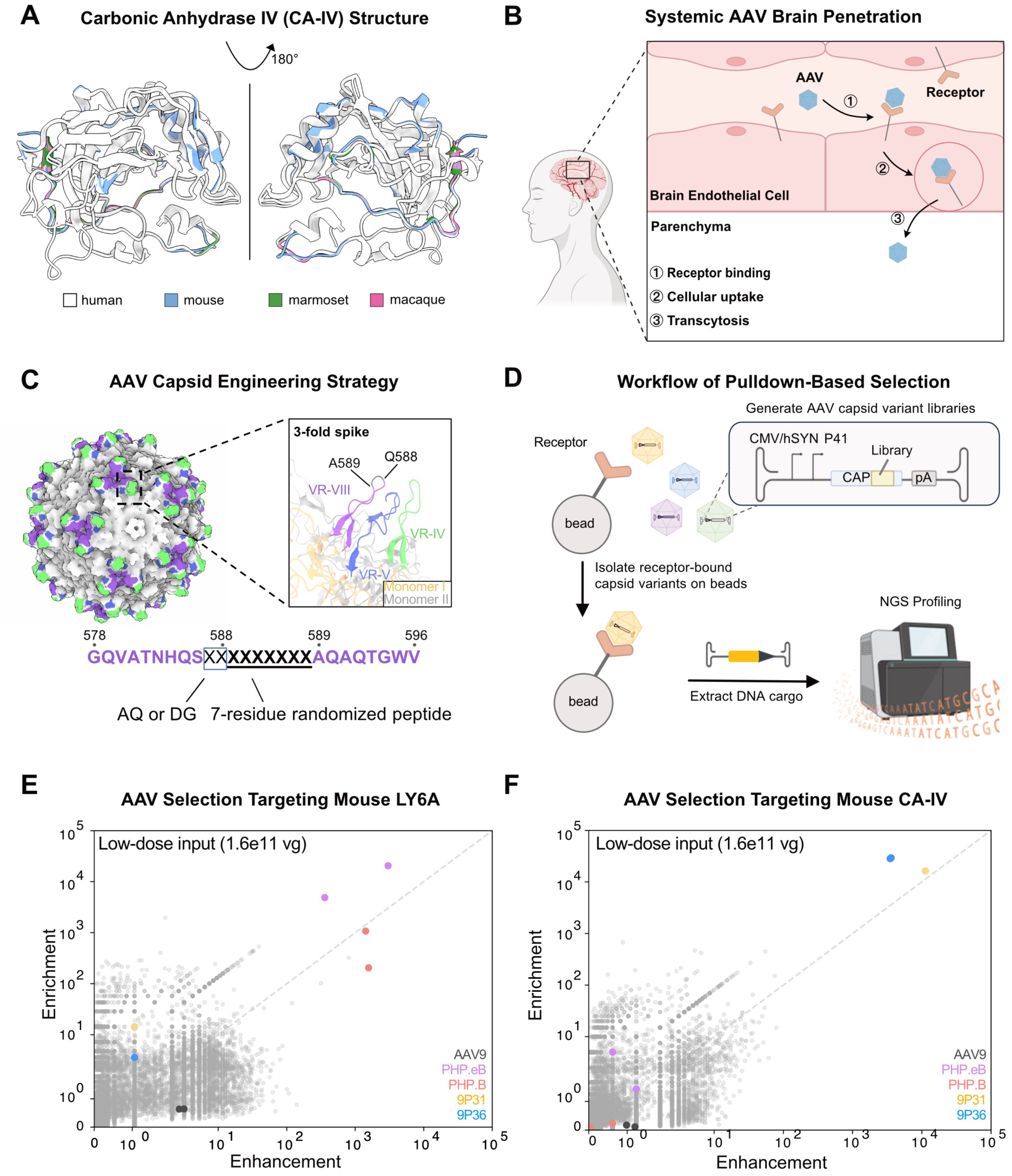
Human blood-brain barrier receptor-targeted AAV capsid engineering pipeline. **(A)** Structure of CA-IV from human (PBD: 1znc^75^), macaque (modeled by Alphafold 3^76^), marmoset (modeled by Alphafold 3) and mouse (PDB: 2znc^77^). Human CA-IV is shown in white. Regions that differ from the human CA-IV in other species, and which may influence receptor binding by AAV variants, are highlighted in blue (mouse), green (marmoset), and magenta (macaque). **(B)** Illustration of critical stages of systemic AAV penetration into the brain from the bloodstream: (1) AAV binding to brain endothelial cell surface receptors; (2) cellular uptake of bound AAV through receptor-mediated internalization; and (3) transcytosis to enter the brain parenchyma. **(C)** Structure of the AAV9 capsid (PDB: 7MT0^77^) highlighting the 3-fold spike and key variable regions (VR) VR-IV (green, residues 445 to 465), VR-V (blue, residues 488 to 511), and VR-VIII (purple, residues 578 to 596). In the enlarged panel, VR-IV and VR-VIII are from the same AAV capsid monomer (colored yellow), while VR-V is from a different monomer (colored light gray). To construct our library of AAV capsid variants, a 7-residue randomized peptide was inserted between positions 588 and 589 of VR-VIII and positions 587 and 588 were modified to either AQ or DG. **(D)** Pulldown-based selection workflow used to identify AAV variants that bind target receptors. Capsid variants that bind to the receptor (e.g. CA-IV or LY6A) are isolated with receptor-bound beads, followed by DNA extraction and next-generation sequencing (NGS) to identify the enriched variants. **(E) (F)** Validation of pulldown-based selection for AAV variants binding mouse LY6A (E) and CA-IV (F) receptors. AAVs known to bind LY6A (PHP.B and PHP.eB) and mouse CA-IV (9P31 and 9P36) show significantly higher enrichment and enhancement than other AAVs in a pool of ∼18,000 unique variants. If both codon replicates are produced, the AAV variant appears as two distinct dots, with each dot representing one codon replicate.

Here, we implemented an *in-vitro* selection pipeline for receptor binding, similar to that recently used for mouse LY6A^31^ and human TfR1^40^, to enrich human CA-IV-binding AAVs. After narrowing down the pool of engineered AAVs specifically binding human CA-IV, single capsid variants were characterized in a “humanized” mouse model expressing human CA-IV in brain endothelial cells. We found that while the top performing capsid variants from receptor binding selections strongly bind human CA-IV and have enhanced potency on human CA-IV-expressing cells, they poorly transduced the “humanized” mouse brain after intravenous administration, with potencies akin to AAV9. Instead, in vivo selections of the same library in a “humanized” mouse revealed that the strongest brain transducers were variants that performed comparatively poorly in binding selections. One such variant, AAV-hCA4-IV77, resulted over 100-fold higher transgene mRNA levels compared to AAV9 and achieved widespread neuronal and astrocytic transduction (between 20–40% of neurons and 15–45% of astrocytes, depending on the brain region). Moreover, AAV-hCA4-IV77 demonstrated approximately 2-fold greater overall cell transduction than AAV9 in human brain organoids, with a similar enhancement in mature neurons. These results suggest that tuning capsid receptor-binding strength may be required for optimal blood-brain barrier transcytosis and support the therapeutic potential of AAVs targeted to human CA-IV.

## Results

### In vitro selections successfully identify known receptor–AAV pairs

The process of receptor-mediated BBB crossing by AAVs involves three key steps: receptor binding, cellular uptake, and transcytosis (Figure 1B)^31,41–43^. First, AAVs circulating in the bloodstream enter blood vessels in the brain and interact with their target receptors on endothelial cells. This interaction then mediates internalization of the AAVs into the endothelial cells. Finally, AAVs are transported through the cells and released into the brain parenchyma. While the mechanistic details of cellular uptake and transcytosis are poorly understood, mechanism-agnostic directed evolution of AAV variants has led to the identification of several luminal brain endothelial receptors, such as LY6A, LRP6 and CA-IV, that enable efficient AAV delivery to the brain following intravenous injection in model organisms^18,31–33,44^. Leveraging this understanding, we designed a receptor-directed *in vitro* engineering workflow to identify AAV capsid variants that effectively interact with validated BBB transcytosis receptors. Our approach builds upon recent successes targeting other BBB receptors (such as murine LY6A and widely-expressed human TfR1)^31,40^.

We first generated a diversified library of AAV variants by modifying the 3-fold spike of AAV9, a natural serotype widely recognized for its brain transduction after systemic administration (Figure 1C). The 3-fold spike is formed by three variable regions: VR-IV, VR-V, and VR-VIII. Applying well-established library design strategies^17,18^, we inserted a randomized 7-amino acid peptide between residues 588 and 589 in the VR-VIII region, combined with either AQ or DG at positions 587-588^45^. As in our previously described M-CREATE method^17^, a low-efficiency transfection of the resulting plasmids into AAV producer cells was performed to maximize the fraction of cells containing instructions for a single library variant and thereby minimize assembly of capsid variant chimeras.

We then used a pulldown-based selection method to identify AAV variants within the library that robustly bind receptors (Figure 1D). We validated this approach using a library which included known receptor-binding AAVs spiked into a background of ∼18,000 unique AAV capsid variants identified through a prior selection to have AAV9-like HEK cell transduction. Each AAV variant was included twice, with synonymous codon replicates of the same protein sequence, to assess reproducibility of subsequent results (sequences shown in supplementary table 1). Mouse receptors LY6A and CA-IV were modified by replacing their C-terminal GPI anchors with an HA tag. We then incubated the AAV library pool with anti-HA magnetic resin containing each HA-tagged receptor. Following an overnight incubation (compare to the 3 to 4 weeks required for *in vivo* directed evolution), we washed and eluted interacting capsid variants. To identify non-specific binders, we performed the same protocol in parallel with anti-HA magnetic resin lacking any receptor.

We used next-generation sequencing (NGS) to evaluate the performance of AAV variants. To quantify the increase in each variant’s abundance post-selection, indicating interaction with the target receptor, we calculated its Enrichment (Er, Equation 1):

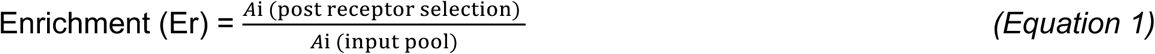

where *Ai* is the relative abundance of variant *i*, calculated as the percentage of variant *i* in the total pool.

To account for non-specific binding to the resin, we also evaluated each variant’s post-selection abundance with and without receptor by calculating its Enhancement (Eh, Equation 2):

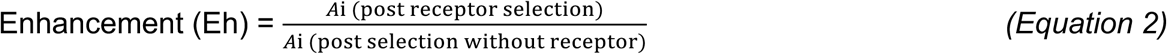

Combined, these metrics provide an overall assessment of binding efficiency and specificity. We noticed one variant, Alpha 1 (9-mer sequence AQYPPVFKS), that showed promising but inconsistent scores across codon replicates in a selection against mouse LY6A. The same variant appeared with low Eh scores consistent with non-specific binding but appreciable Er scores in a selection against mouse CA-IV (Figure S1D, S1E). It is known that LY6A is functional in C57BL/6J and DBA/2J strains of mice but not NOD/ShiLtJ^34^, and that CA-IV is functional in all three strains^32^. We therefore tested the tropism of Alpha 1 in these strains. Interestingly, Alpha 1 showed weakly enhanced endothelial tropism only in NOD/ShiLtJ mice (Figure S1F, S1G), inconsistent with the pattern of either receptor.

As expected, though, AAVs PHP.B and PHP.eB showed high Er and Eh values in selections with LY6A (Figure 1E). Their performance was consistent across both low (1.6 × 10^11^ viral genomes, vg) and high (8 × 10^1^ vg) doses of library input (Figure S1D). Similarly, mouse CA-IV effectively selected AAVs 9P31 and 9P36 (Figure 1F, S1E). The successful identification of known AAV-receptor interactions increased confidence in our selection method.

### In vitro identification of AAVs binding human CA-IV

Having validated our method with known receptor-AAV pairs, we next turned to the identification of novel capsid variants that engage previously-untargeted receptors, specifically human CA-IV. Following the same approach used above for LY6A and mouse CA-IV and previously^31,40^ we conducted one round of selection with our library of AAV9 variants with 9-residue modifications (randomized 7 amino acid peptide insertion between residues 588 and 589 in the VR-VIII region^17^, combined with either AQ or DG substitutions at positions 587-588). By selecting variants showing both enrichment and enhancement scores greater than 3, we efficiently narrowed the vast initial pool down to approximately 10,000 human CA-IV binders.

To prioritize candidate AAV variants for individual validation, we first focused on the top performing capsids during receptor binding selections with human CA-IV, as previous studies demonstrated that AAVs with high enrichment for binding of receptors such as TfR1, LY6A, and LY6C1 exhibit efficient BBB crossing^31,40^. Our experiments also confirmed this pattern, showing that the efficient mouse brain-transducing variant PHP.eB is strongly enriched and enhanced in LY6A binding selections, while 9P31 performs similarly for mouse CA-IV. To evaluate the relative binding strength of capsid variants to human CA-IV, we performed a second round of pulldown-based selection. We produced a vector library of the top 10,000 candidate CA-IV binders along with known AAVs (such as AAV9, PHP.eB and 9P31) as controls, with each capsid variant represented by two synonymous codon replicates (Figure 2A). This selection identified a distinct group of strongly performing human CA-IV binders characterized by high enrichment (Er>15) and enhancement (Eh>40) scores (Figure 2A). The robustness of these results was confirmed by the strong correlation of Er and Eh scores observed between the codon replicates of these AAVs (Figure S2A).

**Figure 2:**
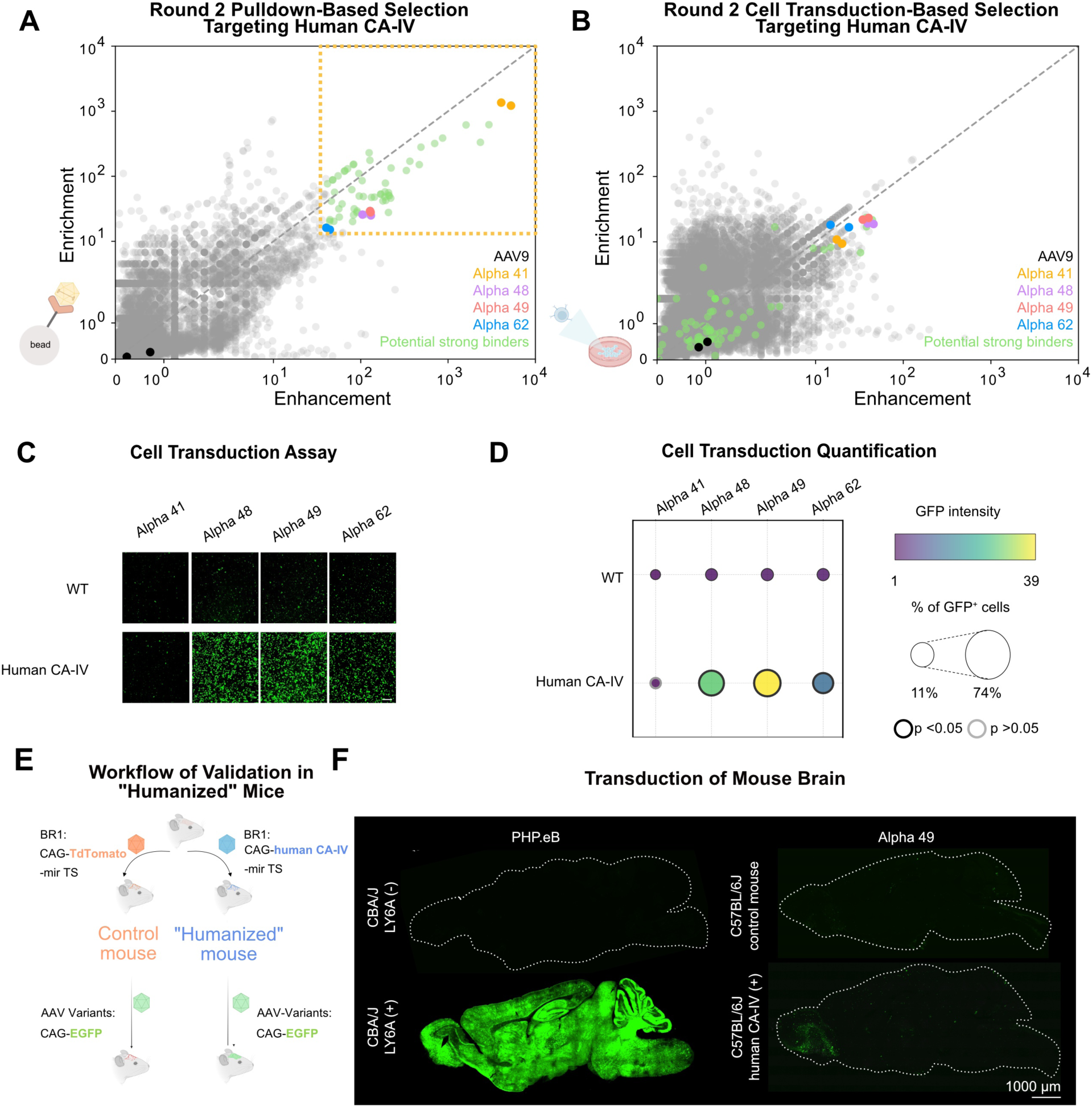
Identification of human CA-IV Targeting AAVs *in vitro.* (A) Results of round 2 pulldown-based selection for AAV variants binding human CA-IV. Promising candidates are highlighted, including the overall top-performing variants (Alpha 41, 48, 49, and 62), along with control AAVs (AAV9) for comparison. Potential strong binders are variants with both high enrichment (Er) and enhancement (Eh) scores (Er>15, Eh>40, top right). (B) Results of round 2 cell transduction-based selection for AAV variants targeting human CA-IV. Promising candidates are highlighted, including the overall top-performing variants (Alpha 41, 48, 49, and 62), along with control AAVs (AAV9). Potential strong binders identified from round 2 pulldown-based selection (2A) are plotted as well. (C) Transduction of wild-type (WT) and human CA-IV-expressing HEK293 cells by overall top performing AAV variants carrying EGFP. Fluorescence images show significant enhancement of transduction by human CA-IV for all variants except Alpha 41. Scale bar, 100 μm. (D) Quantification of cell transduction (EGFP intensity and percentage of EGFP-positive cells), confirming enhanced transduction for Alpha 48, 49 and 62 in the presence of human CA-IV. An independent sample t-test was performed between WT and human CA-IV expressing cells for each AAV variant. Dots with black thick borders represent a p-value of less than 0.05 in both EGFP intensity and the percentage of EGFP-positive cells. Other dots with gray borders represent results that did not meet this significance threshold. (E) Workflow for validation of AAV variants in “humanized” mice. First, 1.5 × 10^12^ v.g. BR1 carrying either CAG-human CA-IV or CAG-TdTomato (control) with microRNA-122 targeting sites (miR-TS, for reducing liver expression) was delivered to establish expression of human CA-IV in brain endothelial cells. After four weeks, 5 × 10^11^ v.g. AAV variants carrying EGFP transgene were administered to evaluate their performance in vivo after three weeks. (F) Representative images of brain sections comparing PHP.eB and Alpha 49 transduction in control mice versus mice expressing human CA-IV or LY6A. PHP.eB demonstrates robust LY6A-dependent brain transduction, validating the model system for evaluating receptor-dependent AAV variants. Scale bar, 1000 μm.

Not all receptor-binding geometries may be compatible with cell internalization (Figure 1B, step 2). To further narrow down the list of high-performing CA-IV binders, we conducted an additional round of screening focused on cell transduction, using the same input pool (∼10,000 AAV variants) as for the round 2 pulldown selection. We transduced HEK293 cells transfected with either human CA-IV or a plasmid containing a scrambled sequence as a negative control and identified AAV variants with CA-IV-boosted transduction through RNA sequencing (Figure S2B). We calculated Er and Eh scores similarly to the pulldown selection, comparing variant abundance in human CA-IV-expressing cells to that in wild-type cells (Figure 2B). The separation between high- and low-performing variants was less than in pulldown-based selections, underscoring the importance of a first round of pulldown-based selections for weeding out non-target-binding AAVs. Integrating our pulldown and cell transduction results yielded a refined list of four candidate variants for further testing: Alpha 41, 48, 49 and 62 (Figure S2C).

Individual characterization by surface plasmon resonance (SPR) with Fc-tagged human CA-IV attached to the chips revealed that all selected variants interact with human CA-IV (Figure S2D). While high avidity precludes accurate affinity determination, results were consistent with sub-nM apparent affinities. We also individually confirmed that identified variant capsids internalize into and transduce HEK293 cells in a human CA-IV-dependent manner. Indeed, human CA-IV binders Alpha 48, 49 and 62, but not Alpha 41, demonstrated CA-IV-enhanced transduction (Figure 2C-D), although the degree of receptor-mediated enhancement was lower for Alpha 62 than for the other two capsid variants.

### A “humanized” mouse model shows that top in vitro CA-IV-binding AAVs fail to transduce the brain

Transcytosis is a necessary third step for BBB crossing (Figure 1B, step 3). Importantly, not all internalized AAVs complete this process. For example, AAV9-X1.1 targets LRP6 across species but primarily transduces endothelial cells rather than crossing the BBB in rodents, while in NHPs the same AAV binding the same receptor results in effective transcytosis and neuronal targeting^33,46^.

To confirm transcytosis *in vivo*, we developed a “humanized” mouse model to investigate BBB crossing. We engineered C57BL/6J mice to express human CA-IV in their brain endothelial cells by systemically delivering either CAG-human CA-IV or CAG-TdTomato (negative control) using a brain endothelial-targeted AAV variant. To prevent potential liver toxicity^47,48^, we incorporated a microRNA-122 target site (mir122-TS) into the cargo construct, which effectively reduces translation in hepatic tissue^46,49^. After four weeks, we systemically administered our AAV candidates carrying the EGFP transgene and evaluated their *in vivo* performance three weeks later (Figure 2E).

To avoid the impact of neutralizing antibodies, which can significantly affect the efficacy of a second dose of AAV^50^, during these sequential AAV administrations, we paired our AAV9-based capsids with the AAV2-based brain endothelial-targeted AAV-BR1^50^. The evolutionary distance between these serotypes has previously been shown to enable efficient brain transduction after sequential administration^50^. As a positive control, we intravenously administered 1.5 × 10^12^ viral genomes (v.g.) of AAV-BR1 carrying either TdTomato (control) or LY6A to CBA mice, a strain known to lack the functional LY6A required for efficient PHP.eB brain transduction^34,35^. Four weeks later, we similarly dosed the mice with 3 × 10^11^ v.g. of PHP.eB CAG-EGFP. As expected, after an additional three weeks, PHP.eB showed a strong dependency on LY6A for effective brain transduction. Only CBA mice engineered to express LY6A in their brain endothelial cells displayed robust EGFP expression across brain regions, while LY6A-deficient TdTomato-expressing controls exhibited minimal transduction (Figure 2F).

Following this validation of our model system, we evaluated the *in vivo* transcytosis efficiency of our engineered AAV variants Alpha 48, 49 and 62. We confirmed expression of human CA-IV in brain endothelial cells, with distribution throughout the cortex, cerebellum, brainstem, thalamus, and hypothalamus (Figure S3A). Sparse neuronal transduction was also apparent in some brain regions, such as cortex and thalamus. Surprisingly, however, we observed poor transduction of our top candidate human CA-IV binding AAVs in both control and “humanized” mice expressing human CA-IV (Figure 2F, S3B).

### In vivo selection of human CA-IV-binding AAVs with efficient BBB transcytosis

This unexpected result prompted us to re-evaluate our engineering pipeline. In our *in vitro* selection process, practical constraints limited our ability to individually test numerous AAVs, leading us to prioritize variants with the most enriched and enhanced human CA-IV binding and cell transduction. However, moderate binding affinity has been reported to enhance BBB transcytosis by antibodies^51^.

Therefore, we utilized our “humanized” mouse model to perform an *in vivo* selection of human CA-IV-binding AAVs (Figure 3A). We intravenously administered 3 × 10^11^ vg of the same pool of human CA-IV binding AAVs (∼10,000 variants), along with control AAVs (PHP.eB and 9P31) used in the prior round 2 binding and cell transduction selections. Each vector carried neuronal synapsin (hSyn) promoter-driven transgene to ensure that when extracting RNA, we would selectively identify variants capable of crossing the BBB rather than those merely efficient at transducing brain endothelial cells (similar to the TRACER method^18^). We injected the AAV pool into both negative control mice expressing tdTomato and mice expressing human CA-IV (n=3 per condition). Three weeks post-injection, we analyzed brain-enriched AAV transgene RNA using the same Er and Eh scoring metrics (Figure 3A).

**Figure 3:**
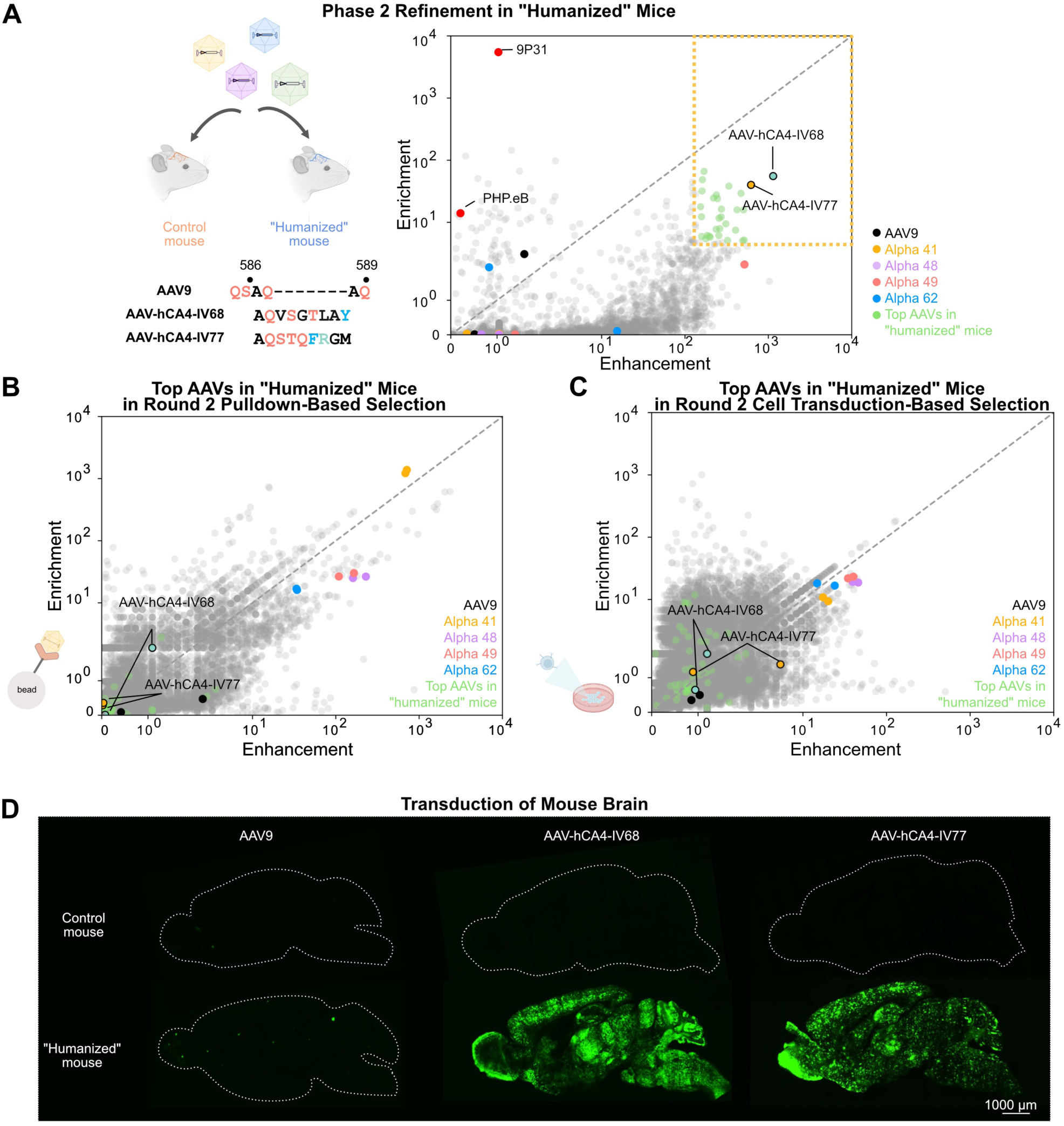
*In vivo* Transcytosis-based selection to filter human CA-IV binders. **(A)** Results of transcytosis-based selection in “humanized” mice expressing human CA-IV in brain endothelial cells. Newly identified variants AAV-hCA4-IV68 and AAV-hCA4-IV77 show high enhancement and enrichment scores, while previously identified strongly enriched and enhanced binders (Alpha 41, 48, 49, and 62) demonstrate low performance in this in vivo context. Control AAVs 9P31 and PHP.eB show high enrichment but low enhancement scores, consistent with their documented brain transduction via mouse CA-IV and LY6A receptors, respectively. Top performing AAVs in “humanized” mice are variants with both high enrichment (Er) and enhancement (Eh) scores (Er>5, Eh>150, colored light green). Amino acid sequences of AAV9 and the newly identified top-performing variants AAV-hCA4-IV68 and AAV-hCA4-IV77. The 9-residue modified regions (positions 586-589) are shown with color-coded based on their properties: black, nonpolar aliphatic; blue, aromatic; red, polar uncharged; green, positively charged; orange, negatively charged. **(B)** Comparison of AAV performance in round 2 pulldown-based selection versus in vivo transcytosis selection. Scores for each variant were averaged across mice (n=3). Notably, the top-performing variants in “humanized” mice (AAV-hCA4-IV68, AAV-hCA4-IV77 and light green colored variants) were not ranked highly in the pulldown selection. **(C)** Comparison of AAV performance in round 2 cell transduction-based selection versus in vivo transcytosis selection. The top-performing variants in “humanized” mice (AAV-hCA4-IV68, AAV-hCA4-IV77 and light green colored variants) were plotted as well. **(D)** Representative brain sagittal sections showing transduction by AAV9, AAV-hCA4-IV68, and AAV-hCA4-IV77 in control mice versus “humanized” mice expressing human CA-IV. To prevent saturation of the fluorescent signal in brain sections, images were captured using reduced illumination intensity. Both AAV-hCA4-IV68 and AAV-hCA4-IV77 demonstrate significantly enhanced and widespread transduction throughout the brains of “humanized” mice compared to control mice, while AAV9 shows consistently poor brain transduction in both groups. Scale bar, 1000 μm.

The control AAVs 9P31 and PHP.eB showed high Er scores and low Eh scores, consistent with their documented robust brain transduction in mice via mouse CA-IV and LY6A receptors, respectively, and independent of human CA-IV. 9P31 demonstrated higher Er than PHP.eB, aligning with previous observations^18^. Among human CA-IV-binding variants, AAV-hCA4-IV68 and AAV-hCA4-IV77 (hereafter abbreviated as IV68 and IV77) outperformed the previously identified strongly enriched and enhanced binders (Alpha 41, 48, 49, and 62), which had low Er and Eh scores in this context (Figure 3A). Interestingly, we noticed that top-performing variants in the “humanized” mouse model (Eh score>150 and Er score>5) were not ranked highly in our earlier round 2 pulldown and cell transduction selections (Figure 3B-C). It is important to note that the Eh and Er scores serve as indicators rather than precise measurements of binding affinities.

When produced as single clones, IV77 exhibited about a 5-fold lower production yield compared to AAV9 and addition of 300 mM NaCl to its buffer helped with stable storage, whereas AAV-hCA4-IV68 achieved yields comparable to AAV9 and required no such buffer formulation. Consistent with their cell transduction selection rankings, whereas Alpha variants showed strong human CA-IV dependent HEK cell transduction (Figure 2C), IV68 and IV77 potency were not affected by human CA-IV expression (Figure S4A-B). Interestingly, only IV77 could be confirmed to have direct binding to human CA-IV by SPR, with an Rmax roughly 30-fold lower than Alpha 49 despite the avidity of both capsid and receptor in the experimental design (Figure S4C). As expected, IV77’s interaction is specific to human and not mouse CA-IV. It is unclear whether the IV68’s lack of binding signal by SPR is the result of exceptionally weak receptor binding or a fortuitous indirect interaction with human CA-IV in vivo.

### Newly identified AAV variants efficiently transduce the brain through human CA-IV

We proceeded with individual validation of IV68 and IV77 in our “humanized” mouse model (Figure 3E). We intravenously administered 5 × 10^11^ vg of AAVs carrying EGFP with a nuclear localization signal (NLS) into both control and “humanized” mice. Three weeks post-injection, we evaluated EGFP signal in brain tissue. The negative control AAV9 showed consistently poor brain transduction in both “humanized” and control mice (Figure 3E). In contrast, both IV68 and IV77 demonstrated significantly enhanced and widespread transduction throughout the brains of “humanized” mice compared to control mice. Unlike PHP.eB and IV68, small patches of particularly strongly-transduced neurons and astrocytes were observed brain-wide after injection with IV77. Analysis across multiple brain regions, including cortex, ventral midbrain, and thalamus, revealed robust transduction by both variants, with neuronal coverage ranging from 20% to 40% and astrocytic coverage from 15% to 45% in human-CA-IV-expressing mice (Figure 4A-B). Notably, GFP signal from both variants was minimal in control mouse brains, requiring higher laser intensity settings to detect their equivalence to AAV9.

**Figure 4:**
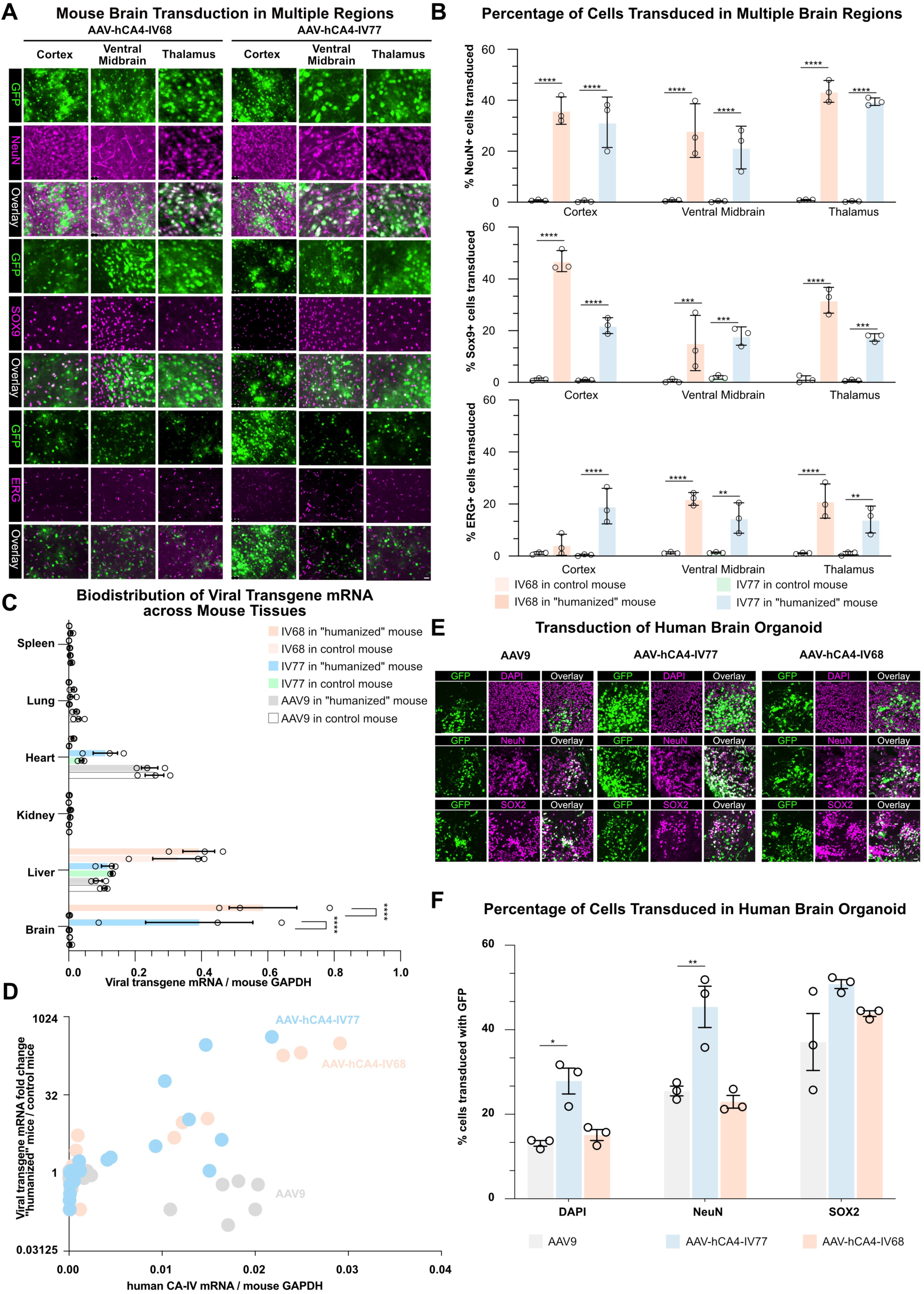
Validation in “humanized” Mouse Model. **(A)** Representative images showing transduction across multiple brain regions (cortex, ventral midbrain, and thalamus) for AAV-hCA4-IV68, AAV-hCA4-IV77, and AAV9 in control and “humanized” mice. Green fluorescence (NLS-EGFP) indicates transduced cells, NeuN marks neurons, SOX9 marks astrocyte and ERG marks endothelial cells. Scale bar, 20 μm. **(B)** Quantification of transduction efficiency across brain regions. The top panel shows the percentage of NeuN+ neurons transduced, the middle panel shows the percentage of SOX9+ astrocytes transduced, and the bottom panel shows the percentage of ERG+ endothelial cells transduced for each AAV variant in control and “humanized” mice. Statistical significance was determined using two-way ANOVA followed by post-hoc multiple comparison tests (Tukey’s test). Asterisks indicate levels of significance (*p<0.05, **p<0.01, ***p<0.001, ****p<0.0001). All experiments were performed in biological triplicate. **(C)** Biodistribution of viral transgene mRNA across multiple organs (brain, liver, kidney, spleen, heart, and lung) in control and “humanized” mice. Data is presented as the ratio of viral transgene mRNA to mouse host genome GAPDH transcript. Statistical significance was determined using two-way ANOVA followed by post-hoc multiple comparison tests (Tukey’s test). Asterisks indicate levels of significance (*p<0.05, **p<0.01, ***p<0.001, ****p<0.0001). All experiments were performed in biological triplicate. **(D)** Correlation between human CA-IV expression level and viral genome distribution across organs. The fold change in viral transgene mRNA levels between “humanized” and control mice is plotted against human CA-IV transcript copies. AAV-hCA4-IV77 (blue circles) and AAV-hCA4-IV68 (orange circles) shows a positive trend with increased transcript levels in organs expressing higher amounts of CA-IV, while AAV9 (gray circles) shows no correlation with CA-IV expression. **(E)** Representative images of human brain organoid transduction comparing AAV9, AAV-hCA4-IV77 and AAV-hCA4-68. Organoids were exposed to AAVs in culture media for two weeks. Green fluorescence (GFP) indicates transduced cells, DAPI marks cell nuclei, NeuN marks mature neurons, SOX2 marks neural progenitors, and overlay shows the co-localization. Scale bar, 20 μm. **(F)** Quantification of transduction efficiency in human brain organoids. Bar graphs show the percentage of total cells (DAPI+), mature neurons (NeuN+), and neural progenitors (SOX2+) transduced by AAV9, AAV-hCA4-IV77 and AAV-hCA4-IV68. Statistical significance was determined using unpaired t-tests. Asterisks indicate levels of significance (*p<0.05, **p<0.01, ***p<0.001). All experiments were performed in biological quintuplicate with individual dots representing independent organoids.

To further characterize these variants, we examined the biodistribution of IV77, IV68 and their dependence on human CA-IV delivered by AAV-BR1, which retains AAV2’s broad peripheral tropism. We quantified viral transgene mRNA across multiple organs, including brain, liver, kidney, spleen, heart, and lung (Figure 4C, S5A-B). As expected, AAV9 cargo transcript levels showed no significant difference between control and “humanized” mice, with predominant expression in liver and heart and relatively low levels in brain. In contrast, IV77 and IV68 demonstrated a clear brain-enriched pattern, reaching levels equivalent to 40% and 60% of the mouse host genome GAPDH transcript in “humanized” mice respectively, while showing minimal expression in control mice. In the brain, both IV77 and IV68 showed over 100-fold higher transgene mRNA levels in “humanized” mice compared to AAV9, and more than 250-fold higher expression compared to its levels in control mice (Figure S5A-B). Additionally, IV68 exhibits slightly increased liver transduction compared to both IV77 and AAV9.

Furthermore, IV77 and IV68 exhibited comparable transcript levels in liver, spleen, lung, and kidney between “humanized” and control mice, with moderately elevated expression in the heart of “humanized” mice. The negligible brain transduction of IV77 and IV68 in control mice also suggests that our engineered AAV specifically recognizes human CA-IV but not mouse CA-IV. This species specificity highlights a key advantage of our selection pipeline—the ability to engineer variants optimized specifically for the intended target species.

To further understand the relationship between human CA-IV expression and transduction efficiency, we investigated whether the transduction patterns of IV77 and IV68 in various organs correlated with human CA-IV expression levels. We plotted the fold change in viral transgene mRNA levels against human CA-IV transcript copies across brain, liver, kidney, spleen, heart, and lung tissues (Figure 4D). While AAV9 showed no correlation with CA-IV expression, IV77 and IV68 demonstrated a positive trend, with increased transcript levels in organs expressing higher amounts of CA-IV. This analysis provides insights into the potential biodistribution profile of IV77 in human therapeutic applications, since human CA-IV expression data is available^37,52^.

As only the BBB was “humanized” in our mouse model, we investigated IV77 and IV68’s potential performance on human brain cells after BBB transcytosis using human brain organoids^53–55^. Human embryonic stem cells (hESCs) were differentiated first into cortical neuron precursors, then aggregated and matured into 3D organoids. AAV variants packaging EGFP under the control of a CAG promoter were added to the culture media (not injected into the organoids), and imaged after two weeks of incubation.

Compared to AAV9, which showed moderate transduction in the organoid, AAV-hCA4-IV77 showed about 2-fold greater overall cell transduction, with a similar improvement in mature neurons (NeuN+) (Figure 4E-F), while IV68 performs similar to AAV9. Since the human CA-IV level in human brain neurons are minimal (Figure S1A), the improved transduction of IV77 may arise through its modulation of the capsid’s naturally evolved receptors^45^.

## Discussion

For the past decade, *in vivo* directed evolution has been the primary method for developing new AAV variants^17,18^. This approach, which can identify capsid variants with enhanced properties without knowledge of the underlying mechanisms, has led to significant advancements in AAV technology. However, it faces challenges in translation due to genetic context differences between species. As directed evolution cannot be performed in humans, there is a significant limitation in the clinical application of engineered AAV capsids identified in this manner. One strategy to minimize this risk is to select AAV variants in multiple species, aiming to identify vectors that utilize BBB crossing mechanisms that are highly conserved; however, these may not be the most efficient in a given species.

Here we utilize a more recent approach to this challenge, engineering optimal AAVs for the intended species by engaging a known BBB transcytosis receptor^31,40^. The choice of luminal BBB receptor is critical to vector potency and safety. Directed evolution of AAVs was greeted with enthusiasm in large part because it offered an alternative to the handful of transcytosis receptors such as TfR1 and INSR that had been known for decades^51,56–58^. As our understanding of AAV transcytosis mechanisms and BBB biology advances^32–35,37,38,52,59–65^, new opportunities for targeted, mechanism-based engineering strategies are emerging. The approach we describe here brings mechanism-guided AAV engineering methods, similar to those recently applied for targeting TfR1^40^, to a promising recently-identified transcytosis receptor utilized by some of the most potent mouse vectors: CA-IV^18^.

Crucially, CA-IV’s expression pattern offers unique benefits over TfR1. Unlike TfR1, which is also expressed in non-brain tissues and non-endothelial brain cells, CA-IV is predominantly restricted to brain endothelial cells (Figure S1A-B)^37,38^. This expression pattern has the potential not only to enhance BBB targeting specificity but also supports future engineering of vectors specific for cell types within the brain. This potential for specificity is supported by the transduction patterns of human CA-IV-binding AAVs in our “humanized” mouse model, where vectors’ peripheral transduction tracks human CA-IV expression, thus providing valuable predictive information for potential biodistribution profiles in human therapeutic applications.

Interestingly, the most effective BBB-crossing variants we identified (AAV-hCA4-IV68 and AAV-hCA4-IV77) were not highly enriched during *in vitro* human CA-IV binding selections. In contrast, strongly enriched binders such as Alpha 41, 48, 49, and 62 performed poorly *in vivo*. This finding expands on recent work suggesting a similar effect with LY6A^66^ and supports the hypothesis made for other therapeutic modalities, such as antibodies, that weak interactions may be optimal for efficient receptor-mediated transcytosis^51^. While both IV68 and IV77 clearly show human CA-IV-dependent brain transduction in vivo, future work will be necessary to understand the precise mechanism of IV68’s enhanced brain potency given its lack of observable direct receptor binding in vitro. Our results with both IV77 and IV68 suggest that stringent round 2 binding enrichment thresholds^31^ may in fact be counterproductive to the identification of potent in vivo vectors. IV77 demonstrated a 100-fold increase in brain transduction efficiency compared to AAV9 in “humanized” mice, and a 250-fold increase compared to control mice, highlighting the vector’s dependence on human CA-IV and the clinical potential of AAVs targeting human CA-IV.

The “humanized” mouse model used here offers a valuable bridge between traditional *in vitro* assays and direct human testing, providing a more physiologically relevant environment for human CA-IV. Notably, our approach of using endothelial cell-specific AAVs to deliver human receptors offers a substantially faster development timeline than traditional transgenics (weeks versus months to years) and greater flexibility to test emerging human receptors. This approach could serve as a template for developing other receptor-targeted therapies, potentially reducing late-stage clinical failures by better predicting human performance during preclinical development.

In summary, here we describe the development of AAV variants specifically engineered for human applications, with a focus on enhanced delivery to the brain by crossing the BBB via the recently-identified receptor CA-IV. Integrating rapid *in vitro* engineering and “humanized” *in vivo* selections, we successfully identified human CA-IV-specific capsid variants capable of efficient transcytosis. Our work addresses the cross-species challenges commonly encountered with murine or NHP-selected AAV variants while highlighting the promise of novel BBB transcytosis receptors identified via those products of directed evolution.

## Acknowledgements

We thank Catherine Oikonomou for help with manuscript editing, Cynthia Arokiaraj for feedback on the manuscript, Xiaozhe Ding for help with AAV structure modeling, Sripriya Ravindra Kumar for guidance on AAV engineering, Nicole B. Yang, Noor E. Ibrahim, Ana S. Jaramillo, Yaping Lei and Zhe Qu for help with AAV production, Richard Hongyi Li for helpful discussions and Zhe Qu, Yaping Lei, Izabela Giriat and Kristen De La Torre for help with AAV evaluation. Figures were made in BioRender. Funding: This project was supported by the Beckman Institute CLOVER Center (to T.F.S. and V.G.), NIH PIONEER DP1NS111369 (to V.G.), Merkin Institute for Translational Research (to V.G. and T.F.S.), NIH BRAIN Initiative Armamentarium UF1MH128336 (to V.G. and T.F.S.), U24MH131054 (to T.F.S. and V.G.), and Friedreich’s Ataxia Research Alliance Postdoc Fellowship, Friedreich’s Ataxia Research Alliance Postdoctoral Research Award (to C.L.).

## Author Contributions

C.L., V.G. and T.F.S. conceived the project. C.L., T.F.S., and V.G. wrote the manuscript and prepared the figures, with contributions from all authors. C.L., J.A.C., and Y.L. produced the AAVs. C.L. and X.C. designed the AAV selection pipeline. C.L., J.D.H. and F.R. analyzed the selection results, and carried out *in vivo* validation. E.E.S. and C.A. conducted cell transduction assays. C.L., Y.L., and J.A.C. contributed to selecting human CA-IV-binding AAVs and individually validating their interactions. T.G. and S.J. purified the CA-IV protein and performed SPR assays. Y.F., J.D.H. and B.K. tested AAVs in human brain organoids. I.T. and A.S. evaluated AAVs in C57BL/6J, DBA/2J, and NOD/ShiLtJ mice.

## Methods

### Plasmids

Sequences encoding AAV capsid variants were cloned into the pUCmini-iCAP plasmid (Addgene ID: 103005). The open reading frames of the CA-IV and LY6A receptors were cloned into the pcDNA3.1 plasmid (ThermoFisher) for production in HEK293 mammalian cells. AAV capsid libraries were constructed using a library backbone containing CMV promoter, P41 promoter and TelN digestion sites, following previously published methods^17,18^. These library plasmids can be obtained upon request from the CLOVER Center at the California Institute of Technology (Caltech).

### Animals

All animal experiments were conducted with approval from the Caltech Animal Care and Use Committee and the California State Polytechnic University Pomona Animal Care and Use Committee, adhering to all applicable ethical guidelines. The mouse strains C57BL/6J (000664), DBA/2J (000671), CBA/J (000656) and non-obese diabetic (NOD)/ShiLtJ (001976) were obtained from The Jackson Laboratory (JAX).

### AAV vector production

AAV packaging and purification were carried out as previously described, with modifications to utilize AAV-MAX suspension cells (ThermoFisher, A51217)^67^. In brief, recombinant AAV was produced by triple transfection of cells in suspension using the VirusGEN AAV Kit with RevIT enhancer (Mirus Bio, MIR 8007) at a molar ratio of transgene: AAV capsid: pHelper = 1:2:0.5, as per the manufacturer’s instructions. The total DNA amount was 2 µg per mL of cells. Virus-producing cells and medium were harvested 72 hours post-transfection. Viral particles were purified using iodixanol step gradient columns followed by ultracentrifugation, as previously detailed^67^. AAV-hCA4-IV77 required a final buffer formulation of DPBS with 0.001% (vol/vol) Pluronic F-68 and 300 mM NaCl for stable storage at 4°C. The purified AAVs were quantified through droplet digital PCR (ddPCR, Bio-Rad) following Addgene’s protocol with modifications. Briefly, AAV samples were serially diluted in nuclease-free water to achieve a suitable concentration for ddPCR analysis. The reaction mixture was prepared using 10 μL of ddPCR Supermix for Probes (Bio-Rad), 1 μL of primers binding to ITR regions (final concentration 900 nM), 5 μL of the diluted AAV sample, 10 μL 2X ddPCR EvaGreen Supermix (Bio-Rad) and nuclease-free water to a final volume of 20 μL. The mixture was then partitioned into droplets using the QX200 Droplet Generator (Bio-Rad). Thermal cycling was performed under the following conditions: 95°C for 10 minutes, followed by 40 cycles of 95°C for 30 seconds and 60°C for 1 minute, and a final hold at 90°C for 5 minutes. After cycling, droplets were read using a QX200 Droplet Reader (Bio-Rad), and the concentration of positive droplets was analyzed using QuantaSoft software (Bio-Rad). The viral genome titer was calculated using the Poisson distribution, accounting for dilution factors and sample volumes.

### BBB receptor single cell transcriptomic data analysis

**Data Acquisition and Processing.** Transcriptomic data were obtained from previous publication^52^, which profiled cellular diversity across the adult human brain using single-nucleus RNA sequencing. The dataset, including raw count matrices and metadata, was downloaded from CELLxGENE^68^. All analyses were conducted using Scanpy^69^, implemented in Python v3.12. **Cell Type Expression Analysis.** To investigate the expression of genes implicated in BBB transcytosis across brain cell types, raw count data were normalized using the normalize_total function in Scanpy and log transformed. Gene expression values were subsequently scaled across all cells prior to visualization. Predefined cell type annotations from the original study were used without modification.

A dotplot was generated using Scanpy’s sc.pl.dotplot function to represent scaled expression levels of selected genes across major brain cell types. **Regional Endothelial Cell Analysis.** To examine the spatial distribution of CA-IV expression across brain regions, endothelial cells were subsetted from the dataset based on the cell type annotations provided by Siletti et al. Average CA4 expression was calculated within each brain region and then mapped onto a modified schematic^52^ using Adobe Illustrator, with expression levels represented as a continuous color gradient.

### Pulldown-based AAV selection

GPI anchors in human and mouse CA-IV and mouse LY6A were identified using PredGPI software. These anchors were removed to enable secretion of the receptor proteins into the culture medium from AAV-MAX HEK293 cells (ThermoFisher, A51217). An HA tag was added to the C-terminus to facilitate protein enrichment on HA resin. The receptor proteins with HA tags were expressed in AAV-MAX cells in suspension, with transfection carried out using the VirusGEN AAV Kit and RevIT enhancer as described above. The cell medium, containing the secreted receptors, was collected and stored at −80°C.

Pulldown-based selection was modified from a previously-published protocol^70,71^. 5 µL of magnetic HA resin slurry (ThermoFisher) was incubated with the AAV library (2×10^10^ to 2×10^11^ v.g.) in 3.5 mL of medium containing the secreted receptors, or in fresh medium, and incubated overnight at 4°C with 0.1% Tween-20 on an end-to-end rotator. After incubation, the resin was pelleted and washed four times with TBS-T (20 mM Tris, pH 7.6, 150 mM NaCl, 0.05% Tween-20). Following the final wash, the resin was spun down to remove any remaining buffer, and 150 μL 1X DNA/RNA Shield (Zymo, R1100-50) was added. Capsids were lysed by heating at 95°C for 10 minutes. Proteinase K (5% v/v) was added after cooling, and samples were incubated at 25°C for 15 minutes to degrade capsid proteins. DNA was extracted using the Quick-DNA/RNA viral kit (Zymo, D7021) following the manufacturer’s protocol. The extracted DNA was then processed to add flow cell adaptors around the diversified peptide region using the following primers:

forward, ACACTCTTTCCCTACACGACGCTCTTCCGATCTACAAGTGGCCACAAACCACCA,

reverse, GTGACTGGAGTTCAGACGTGTGCTCTTCCGATCTCCTTGGTTTTGAACCCAACCGG.

Dual Index primers (NEB, NEBNext Multiplex Oligos for Illumina) were used to add indexes for next-generation sequencing (NGS). The samples were then sent to Novogene or the Caltech Millard and Muriel Jacobs Genetics and Genomics Laboratory for NGS analysis.

### Cell transduction-based AAV selection

HEK293T adherent cells were transfected in 6-well plates with either receptor or scramble plasmids using Lipofectamine 3000 (ThermoFisher) according to the manufacturer’s protocol. Two days post-transfection, an AAV library (1×10^9^ to 1×10^10^ v.g.) was added to the cells. After 24 hours, the cells were pelleted and washed with PBS to remove residual extracellular AAVs. RNA was extracted using the RNA Clean Kit (Zymo) following the manufacturer’s instructions. The extracted RNA was reverse transcribed into cDNA using Maxima H Minus cDNA Synthesis Master Mix (ThermoFisher, M1662). The cDNA was then prepared for NGS, as described above.

### Cell transduction assay of AAV variants

HEK293T adherent cells were transfected in 6-well plates as described above. Two days post-transfection, cells were transferred to 96-well plates at 20% confluency. Simultaneously, AAV variants were added at concentrations of 5×10^8^ or 5×10^9^ v.g. per well. Cells were stained with NucBlue Live ReadyProbes Reagent (ThermoFisher, R37605) and imaged 24 hours post-transduction using a Keyence BZ-X700 microscope. Fluorescence images were analyzed using an established quantification protocol to determine the percentage of transduced cells and the fluorescence intensity per transduced area^32^.

### CA-IV protein expression and purification

CA-IV variants were cloned into a mammalian protein expression vector, with an Fc(human IgG1)-Myc-8xHis tag added to the C-terminus of CA-IV. After sequence verification, the constructs were expressed using the AAV-MAX Production System (Gibco) according to the manufacturer’s instructions. At 96 hours post-transfection, cells were pelleted, and the supernatant was filtered through a 0.5 µM filter. The supernatant was incubated with Ni-NTA agarose (Qiagen) for 2 hours, after which the resin was washed with PBS containing 500 mM NaCl. The protein was eluted from the resin with PBS containing 150 mM imidazole.

### Surface plasmon resonance

All SPR experiments were performed on a Sierra SPR-32 instrument (Bruker). Purified C-terminal Fc-fusions of CA-IV receptors were immobilized via the Fc-tag to a protein A-coupled sensor chip at a concentration of 100 nM in HBS-EP+ buffer (Teknova), reaching a capture level of 1000-1200 response units (RU). For each AAV construct, a 2-fold dilution series of AAV samples (starting at 1000 pM) was injected at a flow rate of 10 µL/min for 240 seconds, followed by a 600-second dissociation phase. A sensor chip regeneration step using 10 mM glycine at pH 1.5 was performed between each cycle. All kinetic measurements were analyzed with double reference subtraction.

### AAV Alpha 1 vector administration, tissue processing, and imaging

Adult C57BL/6J, DBA/2J, and NOD/ShiLtJ mice (n=3), aged 6 to 8 weeks and of both sexes, received intravenous injections of 3×10^11^ v.g. AAV via the retro-orbital sinus. Mice were randomly assigned to groups receiving different AAVs. Following a 21-day incubation period, the animals were euthanized using CO_2_ narcosis and transcardially perfused, first with 10 mL of 0.1 M PBS (pH 7.4), followed by an equal volume of 4% PFA in the same buffer. Subsequently, the organs were extracted and subjected to overnight fixation in 4% PFA at 4°C. After fixation, the tissues were rinsed and preserved in 0.1 M PBS with 0.05% sodium azide, maintained at 4°C. Brain tissue was sectioned into 100 μm slices using a Leica VT100S vibrating blade microtome and imaged using a Nikon C2 confocal microscope and Nikon Elements Software at 20x magnification. Maximum intensity projections of 3 optical sections spaced 2 μm apart were used for quantification using ImageJ and Python.

### AAV selection in “humanized” mouse model

Adult C57BL/6J mice (6-8 weeks old, male, n=3) received intravenous injections of 1.5×10¹² v.g. of AAV-BR1 carrying either CAG-TdTomato or CAG-human CA-IV constructs via the retro-orbital sinus. After four weeks, a second dose of 3×10¹¹ v.g. of AAV pool of top candidates from in vitro selection was administered intravenously. Following a 3-week incubation period, mice were transcardially perfused with chilled PBS treated with 0.1% dimethyl pyrocarbonate (DMPC). Tissues were harvested, and AAV genomic RNA was extracted using TRIzol reagent and the Phasemaker System (ThermoFisher, A33251) according to the manufacturer’s protocol. The extracted RNA was reverse transcribed into cDNA using Maxima H Minus cDNA Synthesis Master Mix (ThermoFisher, M1662). The resulting cDNA was then prepared for NGS, as described above.

### AAV characterization in “humanized” mouse model

To evaluate AAV tropism, we administered candidate AAV constructs individually (n=3 per group per AAV variant). Four weeks following the delivery of tdTomato (control) or human CA-IV (“humanized”), mice were administered individual AAV constructs (RO, 5×10¹¹ v.g.). Animals were subsequently perfused transcardially with chilled PBS treated with 0.1% dimethyl pyrocarbonate (DMPC). Harvested tissues were either post-fixed overnight in 4% PFA or snap-frozen at −80 °C for parallel analyses. Fixed tissues were washed with PBS and sectioned at 100 μm with a vibrating blade microtome (Leica VT 1200S) for histological assessment. To assess cell-type tropism, sections were blocked for 1 hour at room temperature (RT) with PBS containing 10% normal donkey serum and 0.2% Triton X-100. Tissues were then incubated overnight at 4 °C with antibodies for neurons (Ms NeuN; Invitrogen, MA5-33103, 1:500), astrocytes (Rb SOX9; abcam, ab185966, 1:250), and endothelial cells (Rb ERG; Invitrogen, MA5-32036, 1:50). To assess effective delivery of human CA-IV, heat induced epitope retrieval was performed with Tris-based antigen unmasking solution (Vector Laboratories, H-3301-250) for 2 hours at 95 °C. Sections were then blocked and incubated with primary antibody, as described above, with _species_ anti-human CA-IV (ThermoFisher, 13931-1-AP, 1:500) and chicken anti-GFP (Aves Labs, GFP-1020, 1:500). Tissues were subsequently washed 3x with PBS-T and then incubated with fluorophore-conjugated secondary antibodies for two hours at RT. Nuclei were stained with DAPI (1:10,000) and tissues washed 3x with PBS-T and 3x with PBS before mounting with ProLong™ Diamond Antifade Mountant (Invitrogen, P36965). Slides were imaged using a spinning disk confocal microscope (Dragonfly, Andor; Fusion for software control) coupled with an sCMOS camera (Zyla, Andor). Images were acquired with ×10/×25 objectives (Leica) under the same conditions. Quantification of viral transduction was performed with the maximum intensity projection and analyzed using ImageJ and Python.

### Biodistribution of AAV viral transgene mRNA and human CA-IV in mice

Frozen mouse tissues were homogenized and processed using TRIzol reagent and the Phasemaker System (ThermoFisher, A33251) for total RNA extraction. The extracted RNA was reverse transcribed into cDNA as described above. qPCR was performed to assess the biodistribution of viral transcripts using primers specific for EGFP (viral transgene), mouse GAPDH (host reference transcript), and human CA-IV (BR1-delivered transgene in “humanized” mice) with the following primers:

EGFP:

forward, CTATATCATGGCCGACAAGC, reverse, TGTTCTGCTGGTAGTGGTC.

GAPDH:

forward, AAATTCAACGGCACAGTCAA, reverse, CCCATTTGATGTTAGTGGGG.

Human CA-IV:

forward, AGGCACTGTCTAATATCCCC, reverse, GGAAGTAGTGCCTCAGTTTC.

### Transduction of human cortical brain organoids

Human embryonic stem cells (hESCs) were cultured in E8 medium on Vitronectin-coated plates as described previously^72^. Using the ‘dual-SMAD’ induction protocol, hESCs were differentiated into cortical neuron precursors in monolayer culture^73^. The precursors were then dissociated into single cells and 100,000 precursors were placed into each well of V-bottom low-attachment plates to form cortical organoids. Afterward, the organoids were transferred to 10 cm dishes and placed on a shaker for long-term culture and maturation^74^. On day 70, 1×10^10^ v.g. of AAV variants encoding EGFP under control of a CAG promoter were added to the culture medium, with no direct injection into the organoids. Two weeks later, the organoids were cryo-sectioned (thickness 18 μm), fixed with 4% PFA and stained with primary antibody NeuN (Abcam, ab177487), SOX2 (ThermoFisher, 14-9811-82) followed by donkey anti-mouse 647 secondary antibody (Jackson ImmunoResearch, 715-606-151). Fluorescence images were captured using a spinning disk confocal microscope. Quantification was performed with ImageJ and Python.

---

**Figure S1:**
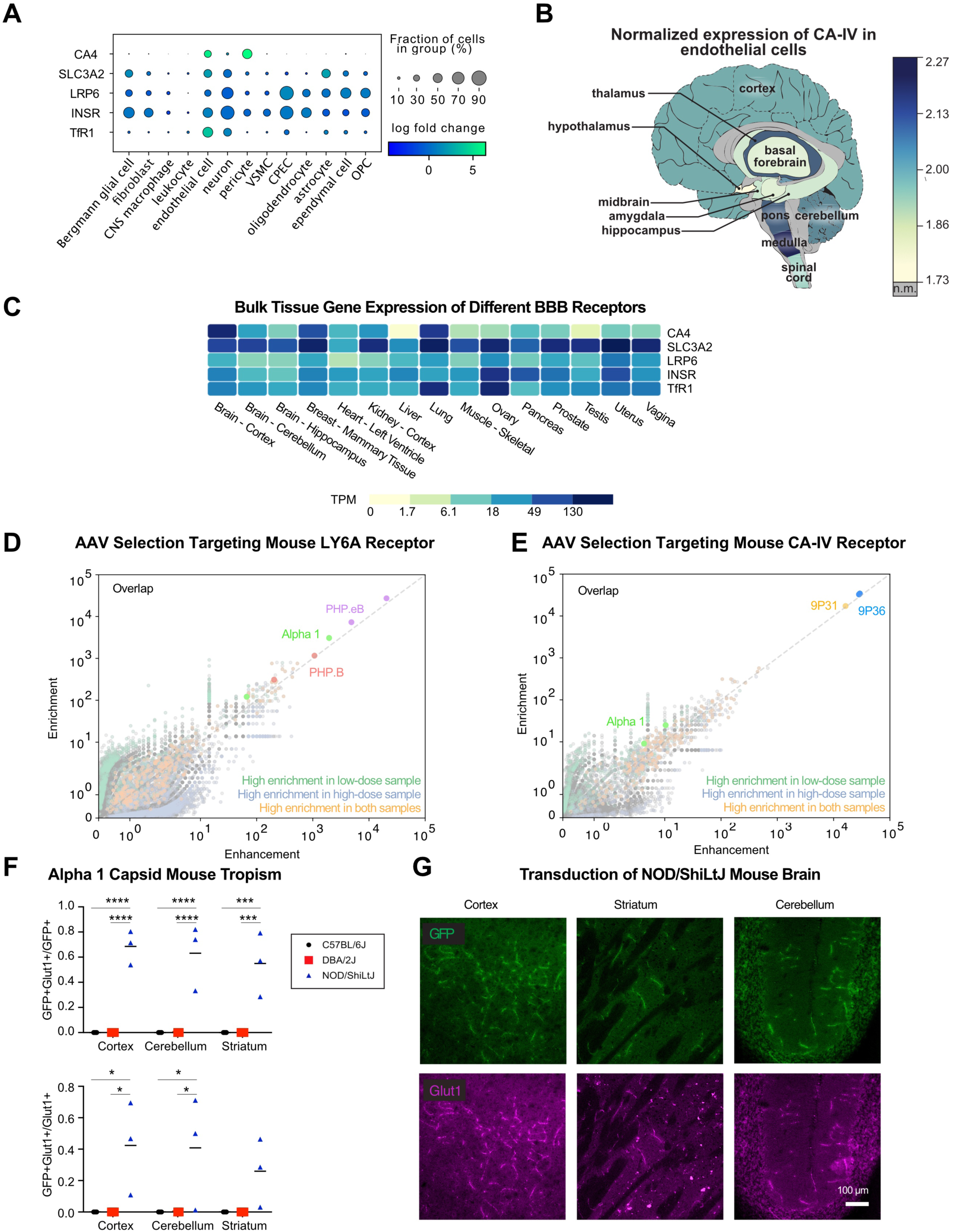
Blood-brain barrier receptor expression profiles and non-specific capsid variant characterization. **(A)** Comparative gene expression profiles of various blood-brain barrier receptors including carbonic anhydrase IV (CA4), solute carrier family 3 member 2 (SLC3A2), low-density lipoprotein receptor-related protein 6 (LRP6), insulin receptor (INSR), and transferrin receptor (TfR1) across different CNS cell types. VSMC, vascular smooth muscle cells. CPEC, choroid plexus epithelial cell. OPC, oligodendrocyte precursor cell. Dot size represents the fraction of cells expressing each receptor (10-90%), while color intensity indicates expression level (log fold change from 0 to 5). CA-IV shows a specific expression pattern across certain CNS cell types compared to other established BBB receptors. **(B)** Heatmap showing the normalized expression of human CA-IV in endothelial cells across different regions of the human brain. CA-IV level in gray colored regions are not measured (n.d.). Data modified from previous publication^52^. **(C)** Heatmap of gene expression levels for several BBB receptors, including CA-IV, solute carrier family 3 member 2 (SLC3A2), low-density lipoprotein receptor-related protein 6 (LRP6), insulin receptor (INSR), and transferrin receptor (TFRC), across various human tissues. Expression levels are quantified as Transcripts Per Million (TPM). Data from GTExPortal^78^. **(D)** Results of selection for AAV variants targeting the mouse LY6A receptor. Comparison of variant performance with low and high input doses reveals consistent enrichment of Alpha 1, PHP.B and PHP.eB. **(E)** Results of selection for AAV variants targeting the mouse CA-IV receptor. Comparison of variant performance with low and high input doses shows reproducible enrichment of Alpha 1, 9P31 and 9P36. **(F)** Quantification of GFP expression in different brain regions (cortex, cerebellum, and striatum) of three mouse strains (C57BL/6J, DBA/2J, and NOD/ShiLtJ) transduced with systemically-delivered Alpha 1 shows tropism inconsistent with known receptors. The top chart shows the percentage of transduced cells that are Glut1+ (endothelial cells), and the bottom the percentage of Glut1+ cells transduced. Animal number =3. Statistical significance was determined using two-way ANOVA followed by post-hoc multiple comparison tests (Tukey’s test). Asterisks indicate levels of significance (*p<0.05, **p<0.01, ***p<0.001, ****p<0.0001). **(G)** Representative fluorescence images of GFP expression (green) in the cortex, striatum, and cerebellum of NOD/ShiLtJ mice three weeks after systemic injection of 3 × 10^11^ vg/mouse of Alpha 1. Colocalization with Glut1 staining (magenta) highlights the transduction of endothelial cells.

**Figure S2:**
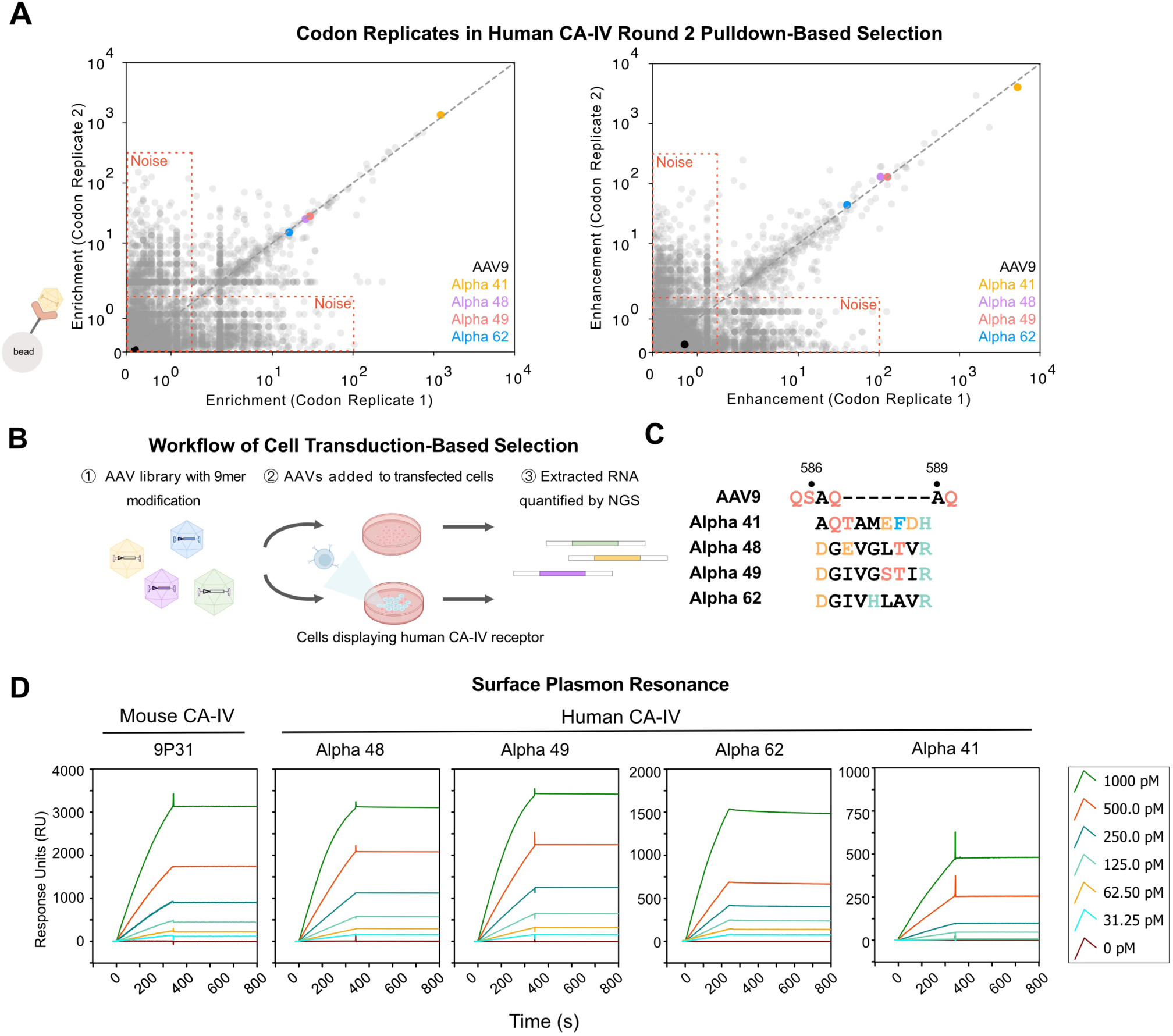
*In vitro* selection and characterization of human CA-IV binders. **(A)** Scatter plots showing the correlation between codon replicates of AAV variants after round 2 pulldown-based selection targeting human CA-IV. High correlation between replicates indicates reproducibility of selection. Variants Alpha 41, 48, 49, and 62 are highlighted. **(B)** Schematic workflow of cell transduction-based selection for identifying AAV variants that effectively transduce cells expressing human CA-IV. The process involves: (1) An AAV library with 9-residue modifications is prepared; (2) AAVs are added to cells transfected with either human CA-IV or a control plasmid; (3) RNA is extracted from transduced cells and quantified by next-generation sequencing (NGS) to identify variants showing receptor-dependent cell transduction. **(C)** Amino acid sequences of the top AAV variants (Alpha 41, 48, 49, and 62) and reference AAV9 capsid showing the 9-residue modified regions at positions 586-589. Amino acids are shown with color-coded based on their properties: black, nonpolar aliphatic; blue, aromatic; red, polar uncharged; green, positively charged; orange, negatively charged. **(D)** Surface plasmon resonance (SPR) binding profiles of AAV variants Alpha 41, 48, 49, and 62 and control (9P31) to mouse and human CA-IV receptors. Response units (RU) are plotted against time (s) at different concentrations (0-1000 pM) of each AAV variant.

**Figure S3:**
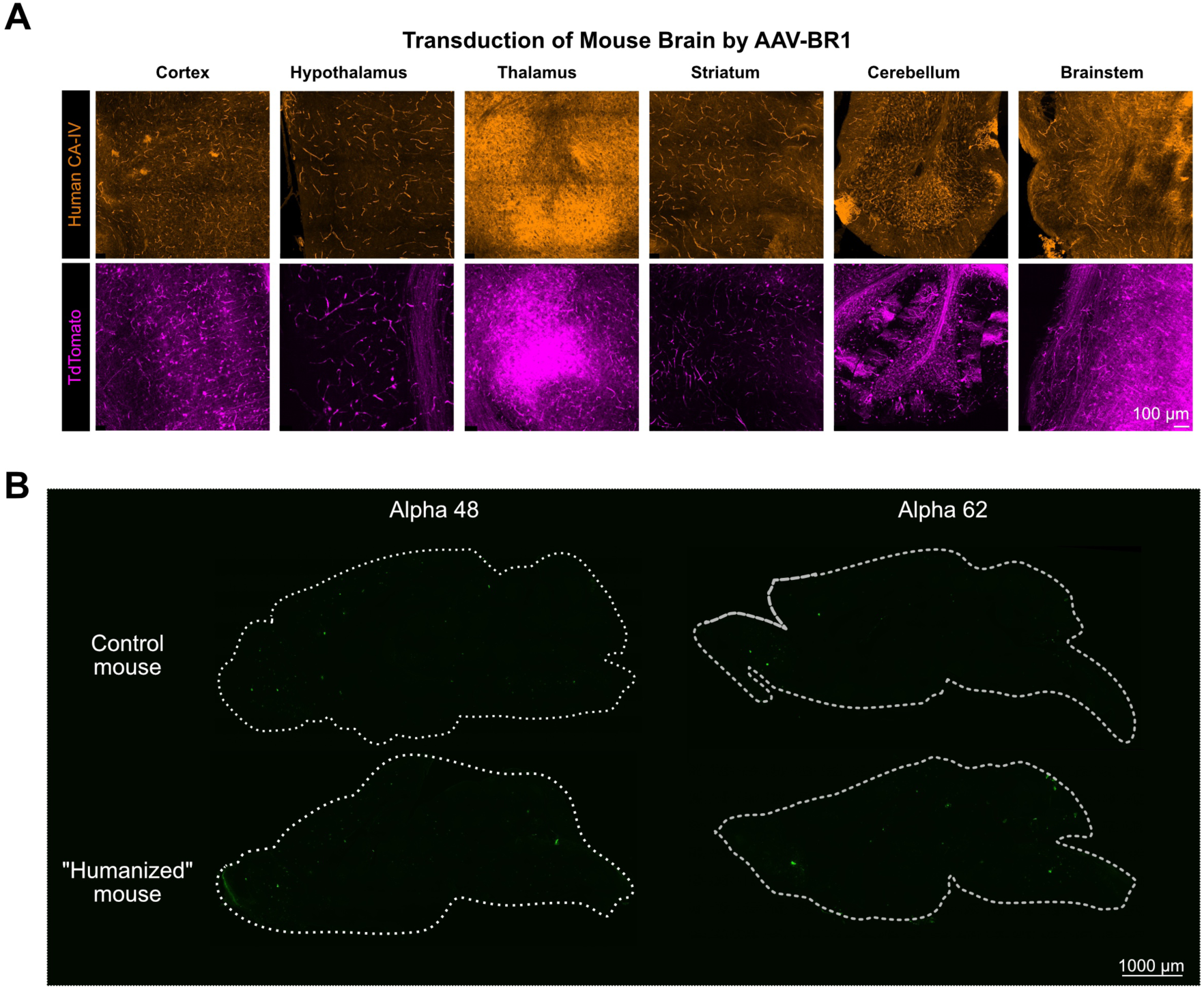
Validation of CA-IV expression in “humanized” mouse and characterization of CA-IV-binding AAVs. **(A)** Detection of protein expression for AAV-BR1 transgenes human CA-IV and TdTomato across multiple brain regions in the “humanized” mouse model. Brown staining indicates immunohistochemical detection of human CA-IV protein and purple staining indicates TdTomato direct fluorescence, which are distributed throughout cortex, hypothalamus, thalamus, striatum, cerebellum, and brainstem confirming successful expression of the human receptor in brain endothelial cells following AAV-BR1 delivery. Scale bar, 20 μm. **(B)** Representative brain section images comparing Alpha 48 and Alpha 62 transduction in control mice versus “humanized” mice expressing human CA-IV. Brain outlines are indicated by dotted white lines. Scale bar, 1000 μm.

**Figure S4:**
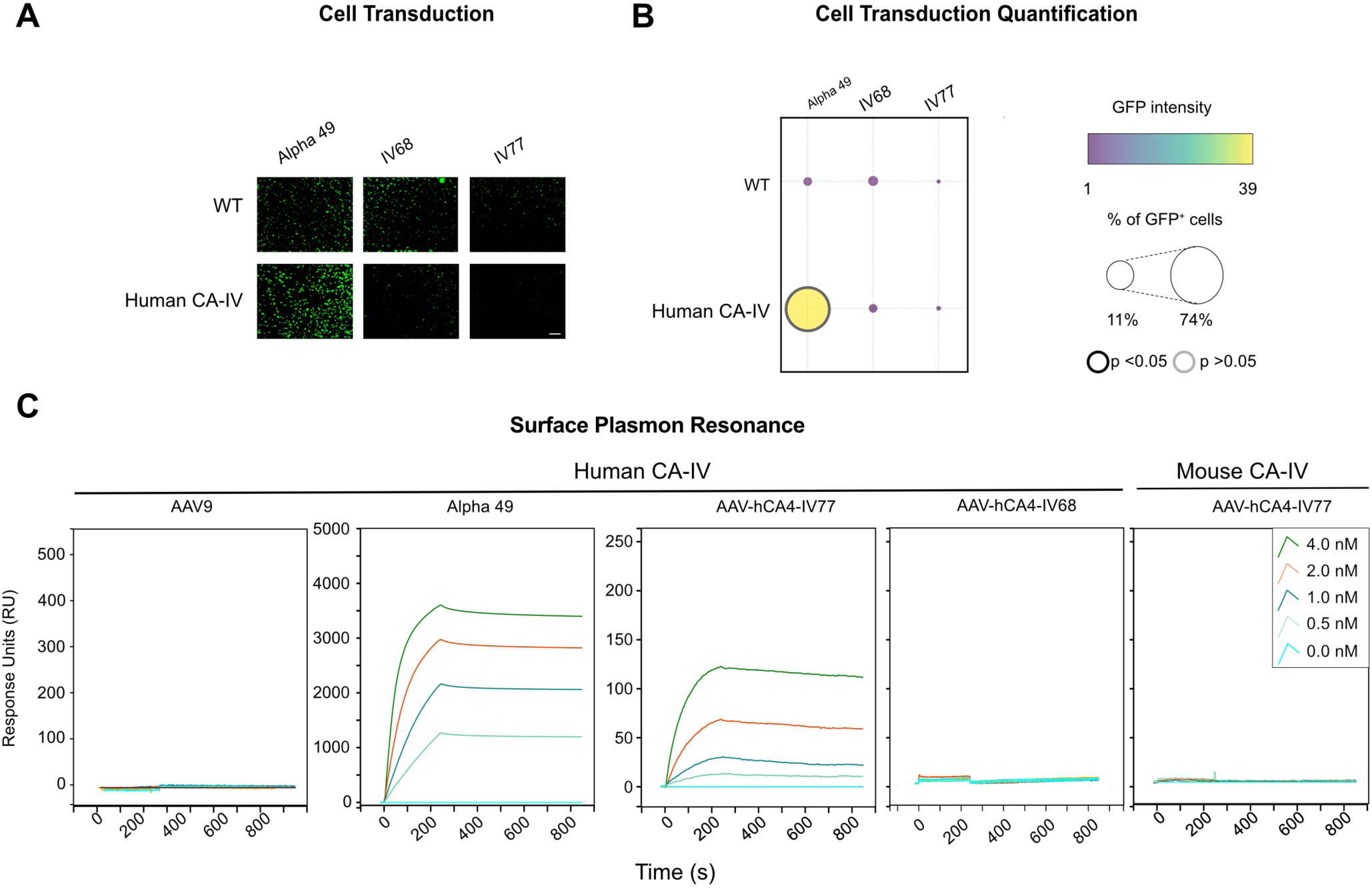
*In vitro* characterization of newly identified AAV variants. **(A)** Transduction of wild-type (WT) and human CA-IV-expressing HEK293 cells by Alpha 49, IV68 and IV77 carrying EGFP. Fluorescence images show only significant enhancement of transduction by human CA-IV for alpha 49. **(B)** Quantification of cell transduction (EGFP intensity and percentage of EGFP-positive cells), confirming enhanced transduction for Alpha 49. An independent sample t-test was performed between WT and human CA-IV expressing cells for each AAV variant. Dots with black thick borders represent a p-value of less than 0.05 in both EGFP intensity and the percentage of EGFP-positive cells. Other dots with gray borders represent results that did not meet this significance threshold. Scale bar, 100 μm. **(C)** Surface plasmon resonance (SPR) binding profiles of AAV variants Alpha 49, AAV9, IV77 and IV68 to mouse or human CA-IV receptors. Response units (RU) are plotted against time (s) at different concentrations (0-4 nM) of each AAV variant.

**Figure S5:**
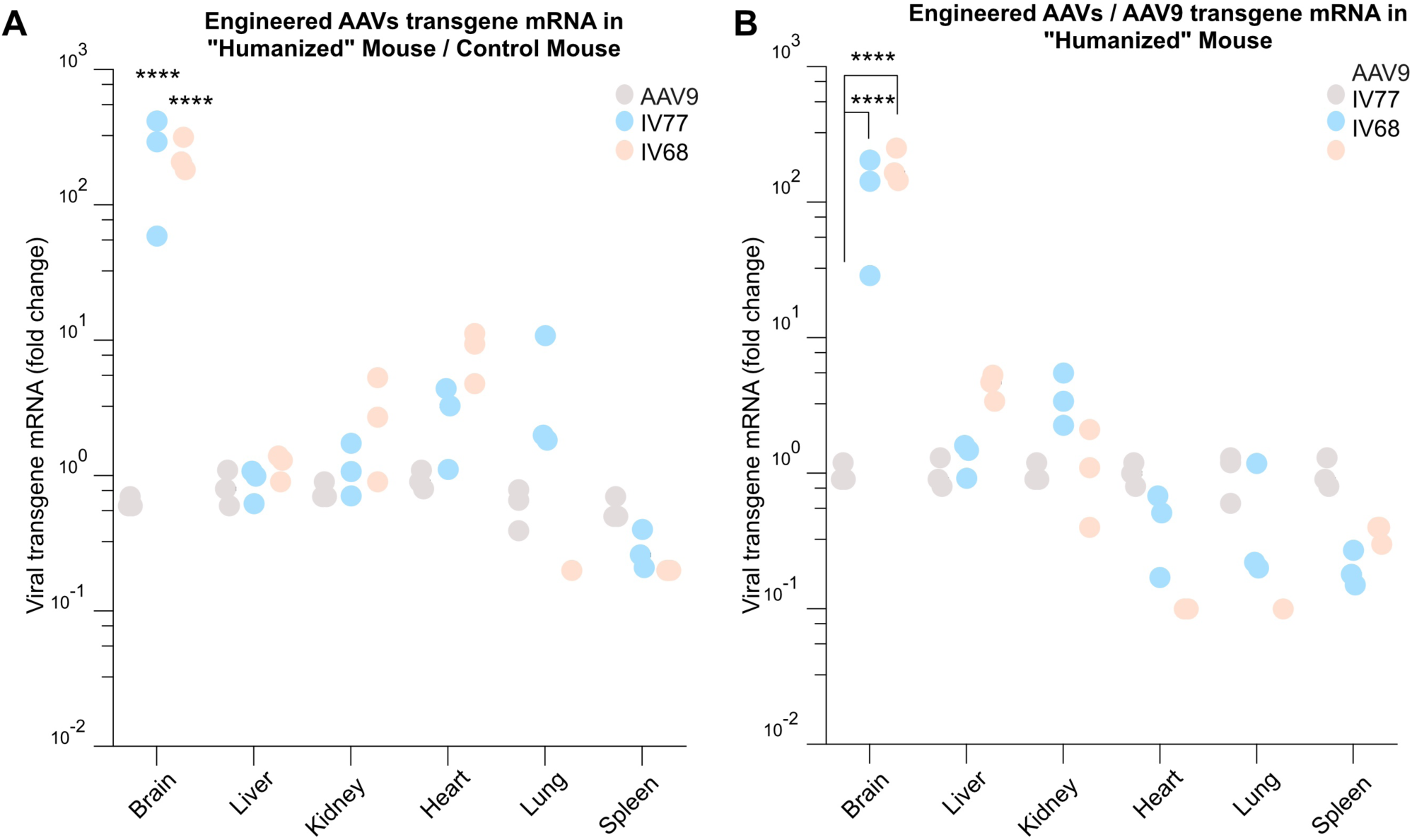
Fold change of AAV transgene mRNA in mouse model. **(A)** Comparison of AAV9, AAV-hCA4-IV77 and AAV-hCA4-IV68 viral transgene mRNA levels across multiple organs (brain, liver, kidney, heart, lung, and spleen) between “humanized” mice expressing human CA-IV (blue circles) and control mice (gray circles). Data is presented as fold change on a logarithmic scale. **(B)** Direct comparison of viral transgene mRNA levels between AAV-hCA4-IV77, AAV-hCA4-IV68 and AAV9 in “humanized” mice expressing human CA-IV. AAV-hCA4-IV77 and AAV-hCA4-IV68 shows significantly higher transcript levels in the brain (>100-fold compared to AAV9), with moderately elevated expression in heart, and comparable or lower levels in liver, kidney, lung, and spleen. Statistical significance was determined using two-way ANOVA followed by post-hoc multiple comparison tests (Tukey’s test). Asterisks indicate levels of significance (*p<0.05, **p<0.01, ***p<0.001, ****p<0.0001). All experiments were performed in biological triplicate.

